# Improving Perennial Ryegrass Transformation Protocols with Developmental Regulators

**DOI:** 10.1101/2025.10.14.682379

**Authors:** Jonathan P Cors, Gabriel W Olson-Jensen, Michael J Smanski, Eric Watkins

## Abstract

The development of genetically modified (GM) or gene edited (GE) turfgrass requires transformation systems that are both efficient and broadly applicable across genotypes. However, traditional *Agrobacterium*-mediated callus culture methods remain limited by low transformation efficiency, extended culture durations, and strong genotype dependence. In this study, we compare a modified classical callus culture protocol with an approach that incorporates the developmental regulator genes *WUSCHEL2* (*wus2*) and *BABY BOOM* (*bbm*), along with an inducible Cre recombinase and a dual luciferase assay to test variable promoter strengths in perennial ryegrass (*Lolium perenne* L.). We show that traditional protocols failed to regenerate plants, despite successful callus formation and transgene expression. In contrast, the developmental regulator system enabled efficient callus induction and plant regeneration independent of genotype. This optimized protocol significantly reduces the time and genotype constraints of perennial ryegrass transformation, offering a practical platform for advanced genetic engineering applications of an important turf and forage grass.

## Introduction

Turfgrass plays a vital role in various landscapes, including recreational areas, sports fields, and urban green spaces. Beyond its aesthetic and functional uses, turfgrass also contributes to environmental sustainability by reducing soil erosion and runoff, and can promote mental wellbeing (Braun et al., 2024). However, turfgrass management presents significant challenges due to its susceptibility to environmental stressors, pathogens, and weed competition (Fan et al., 2020). In recent years, genetic modification has emerged as a powerful tool for plant breeders to target specific improvements in turfgrass, such as introducing genes for enhanced herbicide resistance (Hartman et al., 1994) and stress tolerance (Dai et al., 2003).

Despite the potential advantages of genetically modified (GM) or genetically edited (GE) turfgrass, its widespread adoption is limited by ecological and regulatory concerns. A primary issue is the risk of uncontrolled spread, as GM turfgrass may facilitate gene flow to wild relatives or unintended environments. This concern was exemplified by the escape of a glyphosate-resistant GM creeping bentgrass (*Agrostis stolonifera* L.) from a field trial in Oregon in 2006 (Reichman et al., 2006). To safely release a GM turfgrass, robust biocontainment strategies are essential. Several strategies are currently under development, including apomixis (Wieners et al., 2006), male sterility (Luo et al., 2005), and Engineered Genetic Incompatibility (EGI) (Clark and Maselko, 2020; Maselko et al., 2020; Zinselmeier et al., 2025). However, these approaches are often complex to engineer into a species, and in cases such as EGI, require multiple genetic modifications. A high-efficiency, streamlined transformation protocol is a critical prerequisite for enabling these technologies.

Two main methods have been utilized for turfgrass transformation. Particle bombardment has been used to generate glyphosate-resistant creeping bentgrass (Gardner et al., 2003) and Kentucky bluegrass (*Poa pratensis* L.) (Blume et al., 2010), but suffers from very low efficiency. The second approach, *Agrobacterium*-mediated callus culture, is more commonly used (Singh and Prasad, 2016); this method involves inducing embryonic callus formation using growth hormones, followed by gene transfer via *Agrobacterium*, and then changing the hormones to regenerate roots and shoots from the calluses. However, successful regeneration of viable transgenic plants remains a major hurdle, often limited by genotype specificity and frequent production of albino plants. Furthermore, the callus culture process is time-intensive and prone to multiple failure points, notably from contamination, a key concern in fungal-endophyte containing species such as perennial ryegrass (*Lolium perenne* L.) (Bidabadi and Jain, 2020; Han et al., 2011; Young et al., 2013).

These challenges are not unique to turfgrass. Many monocot species pose similar difficulties in genetic transformation. To address this, N. Wang et al. (2023) demonstrated a novel strategy involving the developmental regulator genes *Wuschel2* (*wus2*) and *Babyboom* (*bbm*), which promote embryogenesis and callus formation with reduced reliance on exogenous hormones. When paired with a heat-shock-inducible Cre recombinase system, these genes can be excised post-induction, allowing for normal regeneration to proceed. This strategy has proven effective in multiple monocot species but has not yet been tested in perennial grasses.

Perennial ryegrass is a cool-season turfgrass commonly cultivated in northern regions of the United States and across Europe. It is also used as a forage crop in agricultural systems (Bonos and Huff, 2015). Between 2010 and 2019, the United States exported 681 million kilograms of perennial ryegrass seed, generating $194 million in revenue (Petty et al., 2024). Several *Agrobacterium*-mediated transformation protocols have been developed for this species, targeting a variety of traits (Bajaj et al., 2006; Grogg et al., 2022; Kumar et al., 2022; Zhang et al., 2020); however, these methods continue to suffer from low efficiency, low throughput, and strong genotype dependence. This genotype specificity is especially problematic given that perennial ryegrass is a cross-pollinated species. Relying on only a few amenable genotypes increases the risk of inbreeding depression and limits breeding progress (Bonos and Huff, 2015).

In this study, we compare the efficacy of traditional *Agrobacterium*-mediated callus culture transformation with the developmental regulator gene approach developed by N. Wang et al. (2023), incorporating *wus2* and *bbm*. We evaluate regeneration rates, promoter performance, and transgene expression to determine the feasibility of this method in perennial ryegrass.

The successful application of this optimized protocol would have far-reaching implications for GM turfgrass. It not only would address a key technical limitation in turfgrass biotechnology, but also enable the implementation of advanced biocontainment strategies. Moreover, the method offers potential for further optimization, increasing its utility for stacking complex genetic traits necessary for next-generation turfgrass varieties.

## Results

### Callus Induction and Transformation Without Developmental Regulators

To assess the feasibility of genotype-independent transformation, we modified methods based on Bajaj et al. (2006) (Figure 1a). A new mercury-free seed sterilization protocol was developed that successfully reduced endophyte and surface contamination. Importantly, we found that fungicide treatment was necessary to eliminate the endophyte (Supplemental Figure 1). While early-stage contamination occasionally occurred, contamination-free tissue transferred beyond the seedling stage remained contamination-free for the entirety of the protocol.

**Figure 1.**
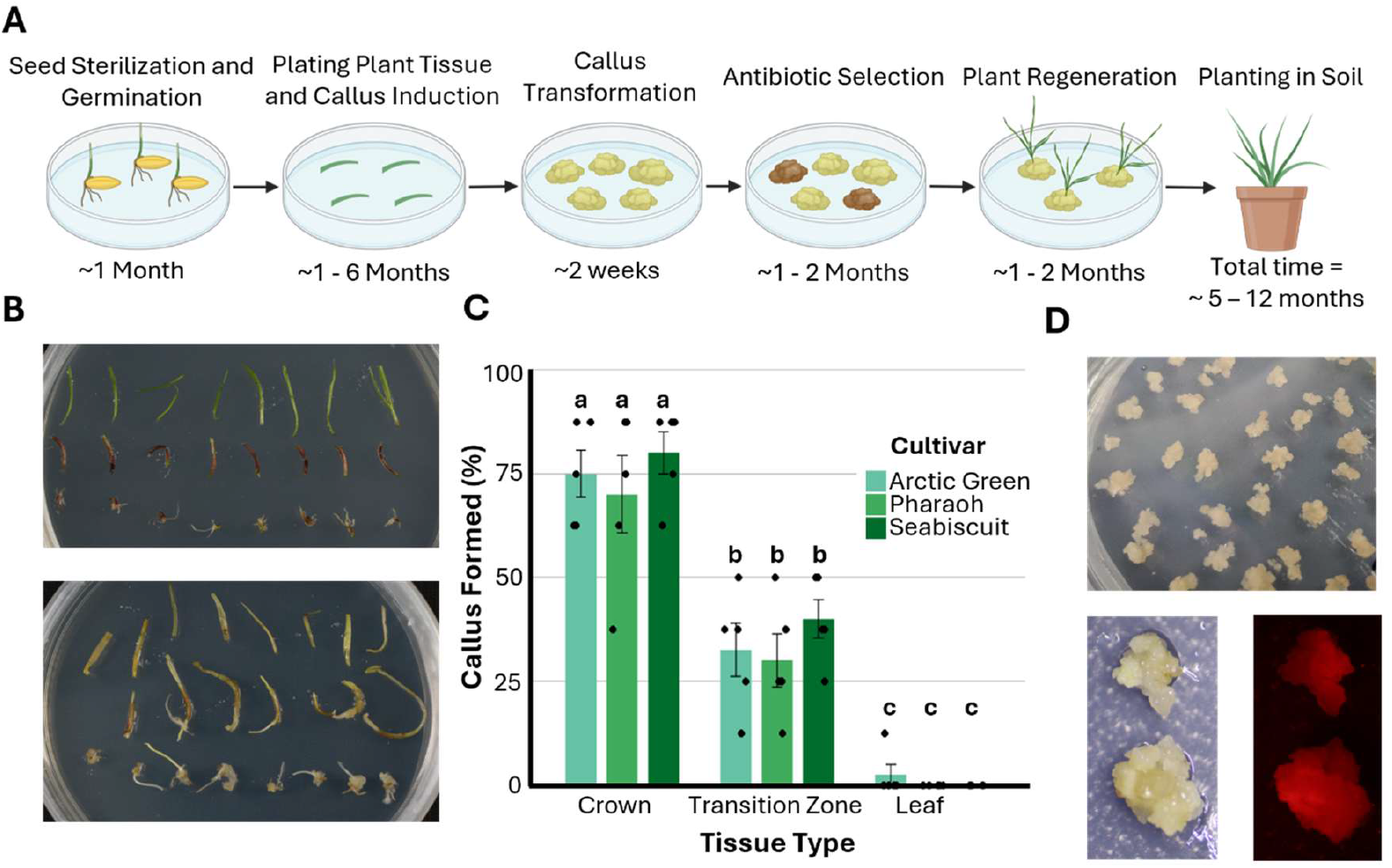
Tissue culture and callus formation efficiency in perennial ryegrass. (A) Schematic overview of the proposed transformation pipeline in perennial ryegrass. (B) Representative images of plated explants before (top) and after (bottom) callus formation. (C) Quantification of callus formation across three tissue types (crown, transition zone, leaf) for three cultivars (‘Arctic Green’, ‘Pharaoh’, ‘Seabiscuit’). Data represent means ± SE (n = 5 biological replicates). Individual data points are shown. Bars sharing the same letter are not significantly different (ANOVA with Tukey’s HSD, *p* < 0.05). (D) Representative images of embryogenic calluses prior to transformation (top), post-transformation (bottom left), and showing red fluorescence (bottom right), indicating successful expression of the reporter gene.

Three tissue types were evaluated for callus induction: crown, transition zone (between crown and leaf), and leaf tissue (Figure 1b). For each of three tested cultivars (‘SeaBiscuit’, ‘Pharaoh’, ‘Arctic Green’), 40 seedlings each were dissected and placed on a callus induction medium. The crown consistently induced the highest callus formation (Arctic Green - 75% ± 5.6, Pharaoh - 70% ± 9.3, SeaBiscuit - 80% ± 5.0), intermediate zones showed moderate induction (Arctic Green - 32.5% ± 6.4, Pharaoh - 30% ± 6.4, SeaBiscuit - 40% ± 4.7), and leaves rarely produced callus, occurring only once in ‘Arctic Green’ (Figure 1c).

Calluses typically formed within one month and could proliferate for up to six months (Figure 1d). Calluses were transformed with a plasmid containing hygromycin resistance and RFP (Supplemental Table 1). Two weeks post-transformation and antibiotic selection, RFP expression confirmed successful transformation in 74% of 104 calluses (Figure 1d, Supplementary Figure 3). However, no transformed callus regenerated into plants.

### Dual Luciferase Assay

A dual luciferase assay was conducted to characterize optimal promoters for transgene expression in perennial ryegrass (Figure 2). The *PvUbi2* promoter demonstrated the highest relative activity, having 5.4 ± 0.25 fold higher expression than CmYLCV. This was followed by the maize and rice ubiquitin promoter, each having 3.55 ± 0.97 and 3.133 ± 0.77 higher fold expression. The promoters Os*EF1a, ZmEf1a, 2x35S* and *Nos* were statistically not different from *CmYLCV*.

**Figure 2.**
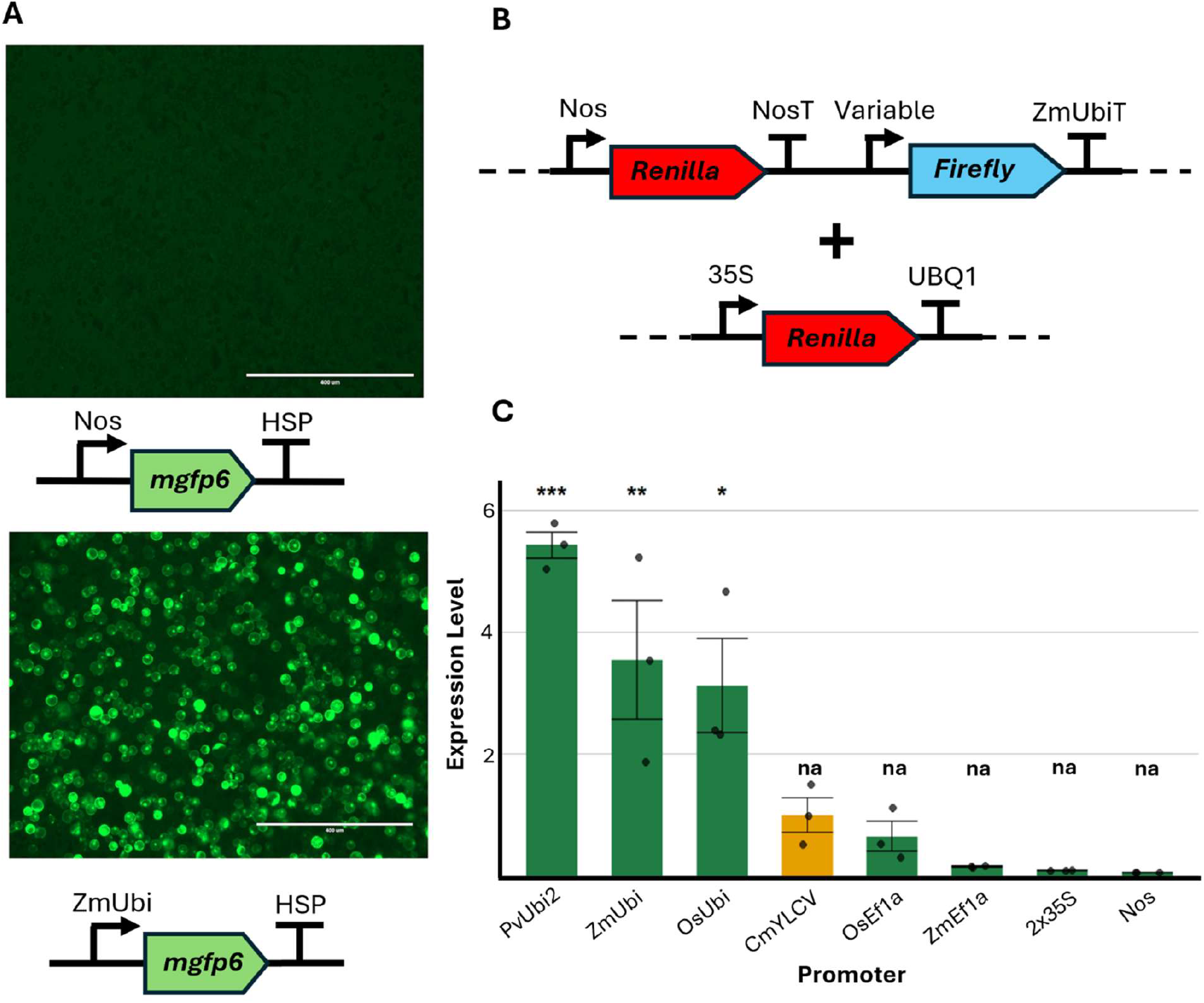
Protoplast transformation and promoter activity in perennial ryegrass. (A) Transient expression of *mgfp6* driven by the Nos (top) or ZmUbi (bottom) promoter in protoplast cells visualized by fluorescence microscopy 24 hours after transformation. Strong fluorescence was observed under the ZmUbi promoter, while little to no signal was detected with the Nos promoter. Scale bars = 400 µm. (B) Schematic of the dual-luciferase reporter constructs used to quantify promoter activity. The experimental construct contains a Firefly luciferase reporter under the control of a variable promoter, and a Renilla luciferase gene driven by a Nos promoter for normalization. A second construct expressing Renilla under the 35S promoter and terminated by UBQ1 was co-transformed due to lack of Renilla signal. (C) Quantification of relative Firefly luciferase activity driven by different promoters in ryegrass protoplasts. Data are normalized to Renilla luciferase and shown as mean ± SD (n = 3). Asterisks indicate statistical significance compared to the CmYLCV promoter (*p* < 0.05, **p** < 0.01, ***p*** < 0.001). “na” indicates expression was not statistically different from CmYLCV.

### Callus Induction & Transformation with Developmental Regulators

To overcome regeneration limitations, we adapted the method of N. Wang et al. (2023) incorporating *wus2* and *bbm* driven by various promoters, with or without *cre* recombinase. Eight constructs were tested in total (Figure 3a, Supplementary Table 1).

**Figure 3.**
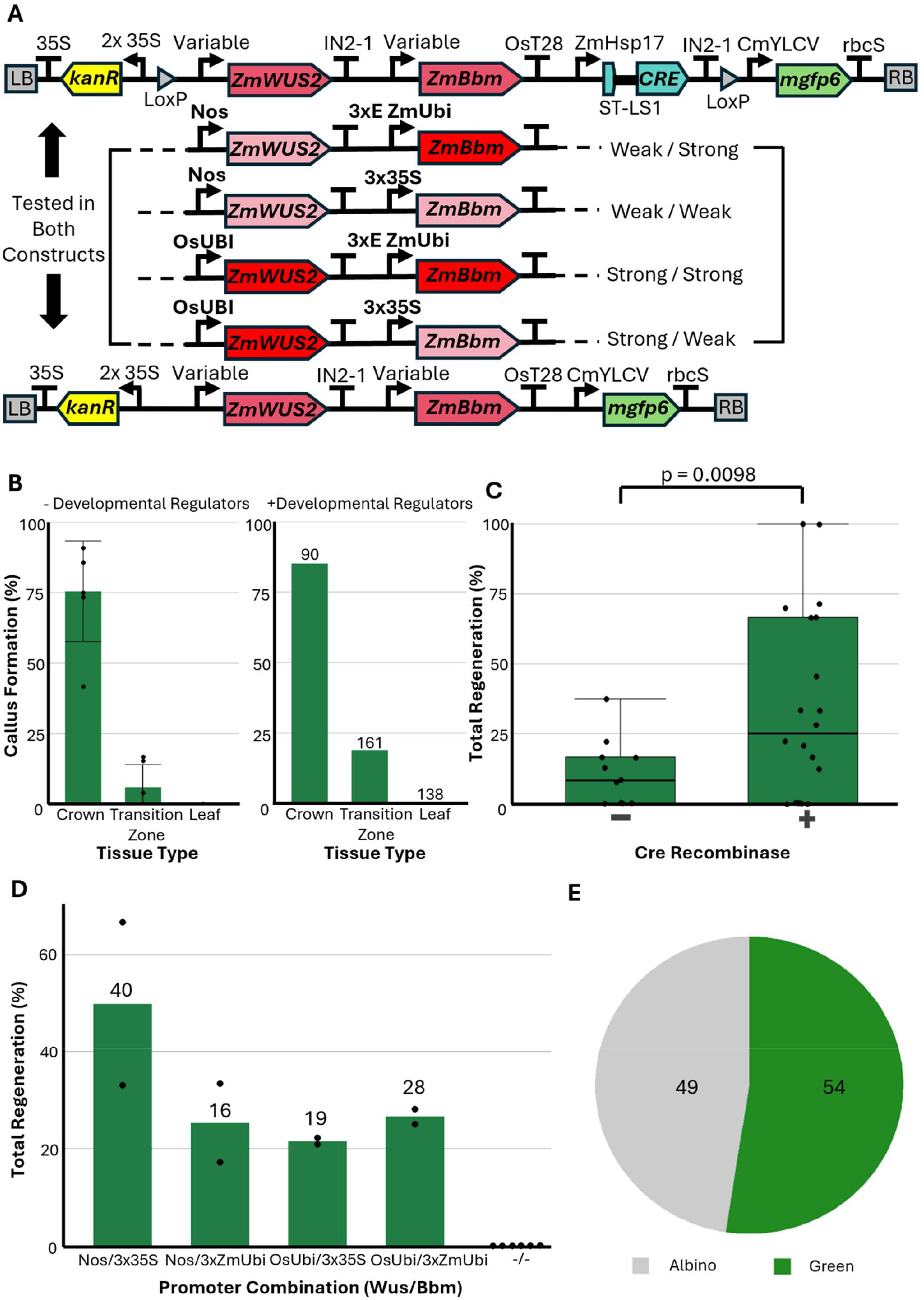
Optimization of regeneration in perennial ryegrass using developmental regulators and Cre-lox recombination. (A) Schematic of transformation constructs containing maize developmental regulators *ZmWUS2* and *ZmBbm* driven by different promoter combinations (*Nos, 3×35S, ZmUbi, OsUbi*), with or without *cre* recombinase and loxP sites. Constructs were tested in parallel to evaluate the impact of promoter strength and Cre-mediated recombination on regeneration. (B) Callus formation efficiency across three tissue types (crown, transition zone, leaf) in the absence (left) or presence (right) of developmental regulators. Numbers above bars indicate total explants per group. (C) Regeneration frequency with or without *cre* recombinase. Addition of *cre* significantly increased total regeneration percentage (*p* = 0.0098, unpaired t-test). (D) Regeneration efficiencies by *wus2/bbm* promoter combination and a negative control without developmental regulators. The number above each bar represents the total number of regenerated plants. (E) Distribution of regenerated plants. Of the total regenerated plants, 54 were green and grew normally, while 49 were albino and did not form roots.

Callus induction was re-evaluated during these experiments and confirmed previous findings that crown tissue yielded the highest callus induction. Intermediate tissue showed reduced rates, and leaf tissue remained non-responsive. This held true whether or not developmental regulators were included (Figure 3b).

Following transformation, tissues were subjected to selection and regeneration. Control transformations lacking developmental regulators failed to regenerate, even when surviving selection. In contrast, tissues transformed with *wus2/bbm* constructs successfully regenerated into plants. The role of *cre* recombinase was tested to assess its impact on regeneration: inclusion of *cre* significantly enhanced regeneration rates changing from a mean regeneration of 11.05 ± 3.6% to 34.4 ± 7.6%, with a p-value of 0.0098 (unpaired t-test) (Figure 3c).

Promoter strength driving the developmental regulators was tested as it is thought to be a key aspect of inducing regeneration. Contrary to this belief, no statistically significant differences were observed between tested promoter combinations. However, this may be caused by having only two replicates and high variability. The promoter combinations *Nos/3x35S, Nos/3xZmubi, OsUbi,3x35S*, and *OsUbi/3xZmubi* were able to regenerate a total of 49.8%, 25.3%, 21.5%, and 26.6%, equating to a total number of plants of 40, 16, 19, and 28, respectively (Figure 3d). Tissue transformed with our negative control plasmid that did not contain the cre and developmental regulators were able to form callus, but none of the callus were able to regenerate. Additionally, of the total 103 plants, 49 were albino and 54 were green and normally growing (figure 3e). There was not a statistically significant difference between promoter strength or inclusion of *cre* in albino and green regeneration.

### Genomic Confirmation of Transformation

Regenerated plants were transferred to the greenhouse (Figure 4ab) and subsequently had DNA extracted from them. From the first and second replicate, 41 and 32 plants were sampled, respectively. In each sample, the expected band was observed matching the positive control while the no template and four different wild type controls each showed no band (Figure 4c, Supplemental Figure 2a). Additionally, a PCR spanning the LoxP joining segment of DNA was performed and it revealed that for 55 out of 73 regenerated plants the developmental regulators were not successfully excised (Figure 4d, Supplemental Figure 2b).

**Figure 4.**
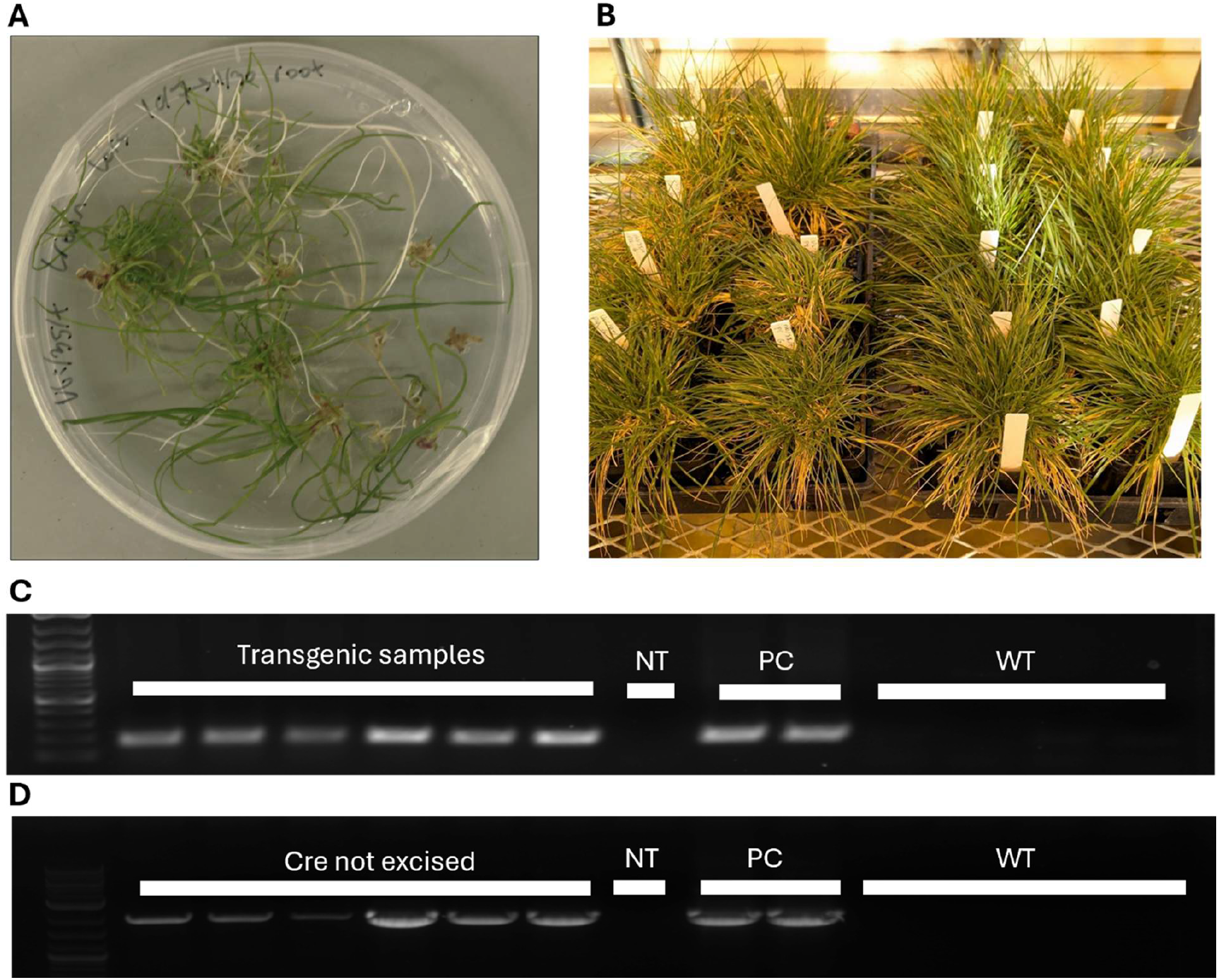
Regeneration and molecular validation of transgenic perennial ryegrass. (A) Regenerated transgenic plantlets prior to soil transfer and (B) established plants growing in the greenhouse. (C) PCR confirmation of transgene presence in regenerated plants. Multiple transgenic samples show amplification of the target band, with no amplification observed in the no template control (NT) or wild-type (WT) samples. Positive control (PC) bands are plasmids Ubi/Zm/+ and nos/35/+. (D) PCR assay detecting Cre-mediated excision of the transgenes.

## Discussion

Developing efficient and broadly applicable genetic transformation protocols is essential for advancing trait engineering in turfgrasses such as *Lolium perenne* (perennial ryegrass). However, transformation in this species remains limited by endophyte contamination, lack of genetic resources, and long arduous protocols with low regeneration rates. Our study addresses these challenges by incorporating developmental regulators into transformation workflows, demonstrating a robust strategy that improves regeneration efficiency and circumvents the need for extensive genotype pre-screening. In doing so, we provide a more accessible platform for generating transgenic ryegrass, with potential applications in both basic research and biotechnological innovation.

A key technical barrier in tissue culture is contamination by endophytes, which are typically beneficial in field conditions as they are able to confer pest resistance (Ma et al., 2025), but can overgrow in sterile environments. To address this, we developed a revised seed sterilization protocol incorporating fungicide treatment. Further improvements could be made to reduce initial contamination observed on seed plates, but plates which were contamination free would remain contamination free for the entirety of the protocol. Additionally, while the removal of endophyte is necessary for transformation, it will likely be beneficial to re-inoculate (Santhanam et al., 2015) the plants after transgenesis to reintroduce positive endophyte effects.

Independent of the cultivar or transformation method, crown tissue consistently produced the highest rates of callus formation. This was followed by the intermediate tissue between crown and leaf, and leaf tissue showing negligible callus induction. This is in line with previous experiments conducted in rice (Hu et al., 2017) which showcase the farther away you get from the crown, the less likely for callus formation. Although crown tissue offers the best callus generation potential, the intermediate zone may be more suitable for certain experiments, as it allows continued propagation of the source plant. Additionally, other tissues not tested within this work may be suitable such as roots, seeds, and anthers (Ikeuchi et al., 2013; Long et al., 2022).

To gain a greater understanding of promoter activity for the expression of the developmental regulators, a dual luciferase assay was employed. Our dual luciferase assay ranked promoter strengths in perennial ryegrass, with *PvUbi2* and other ubiquitin promoters performing best. While our results confirmed trends observed in previous work by Chamness et al. (2023), the overall fold-change was lower in perennial ryegrass than in rice. This may reflect species-specific differences in promoter activity or relatively higher baseline expression of *CmYLCV* in perennial ryegrass, against which the other promoters were normalized.

Despite extensive experimentation with modified versions of established protocols, including variations in tissue type, hormone concentrations, and antibiotic selection, no viable plant regeneration was observed (data not shown). This suggests that the traditional methods are highly genotype-dependent, a limitation also noted by Bajaj et al. (2006) and Grogg et al. (2022), who both screened for responsive genotypes before proceeding with transformation. In contrast, our adoption of the N. Wang et al. (2023) approach proved effective in generating transgenic plants without first screening for conducive genotypes, removing a large bottleneck in the transformation pipeline. In future work we plan to more rigorously assess these methods in more diverse genotypes and cultivars of perennial ryegrass, as well as other species of turfgrass.

While the adoption of the N. Wang et al. (2023) methods proved to be successful in regenerating transgenic plants, several unexpected results were seen. Within our control callus induction tests, both plants that were transformed with *wus2/bbm* and plants transformed without the developmental regulators formed callus at equal rates. This indicates that the media itself is capable of inducing calluses, and that *wus2* and *bbm* do not increase the efficiency of callus induction. However, despite the ability for callus to be induced, only the plants including *wus2/bbm* resulted in plant regeneration. This suggests that the inclusion of the developmental genes may enhance regeneration, a finding also noted in several other studies (Gordon-Kamm et al., 2019; Lowe et al., 2016; J. Wang et al., 2023).

The presence of *cre* recombinase was statistically correlated with successful regeneration, consistent with findings in N. Wang et al. (2023): however, the mechanism underlying this effect remains unclear. It was initially hypothesized to be attributed to Cre-mediated excision of the developmental regulators, but PCR analysis of the *cre* excision revealed that only 25% of regenerated plants had the genes excised. Other factors, such as heat-shock treatment, may also play a role (Dündar et al., 2024), as the control samples without *cre* were not put under a heat shock. No noticeable phenotypic differences have been noted in our plants retaining the recombinase and developmental regulator genes, but negative phenotypes in plants retaining the genes has been noted in other findings (Ikeuchi et al., 2013). Further crossing of the plants may be necessary to remove the genes, or an improved marker system to better track the excision of *cre*, as showcased in Wu et al. (2025), may be desirable.

In conclusion, our study advances the genetic toolkit for perennial ryegrass by demonstrating an effective transformation strategy that eliminates the need for pre-screening of genotypes ability to regenerate and dramatically reduces the overall time of the protocol (Figure 5). Future improvements utilized in N. Wang et al. (2023) but not tested here, such as automated tissue sectioning and the inclusion of growth-enhancing compounds like ancymidol, could further increase throughput and consistency. Additionally, the swapping of confirmations markers from *mgfp6* to the RUBY marker as utilized in Wu et al. (2025) could allow for simpler tracking of transformation and successful *cre* excision.

**Figure 5:**
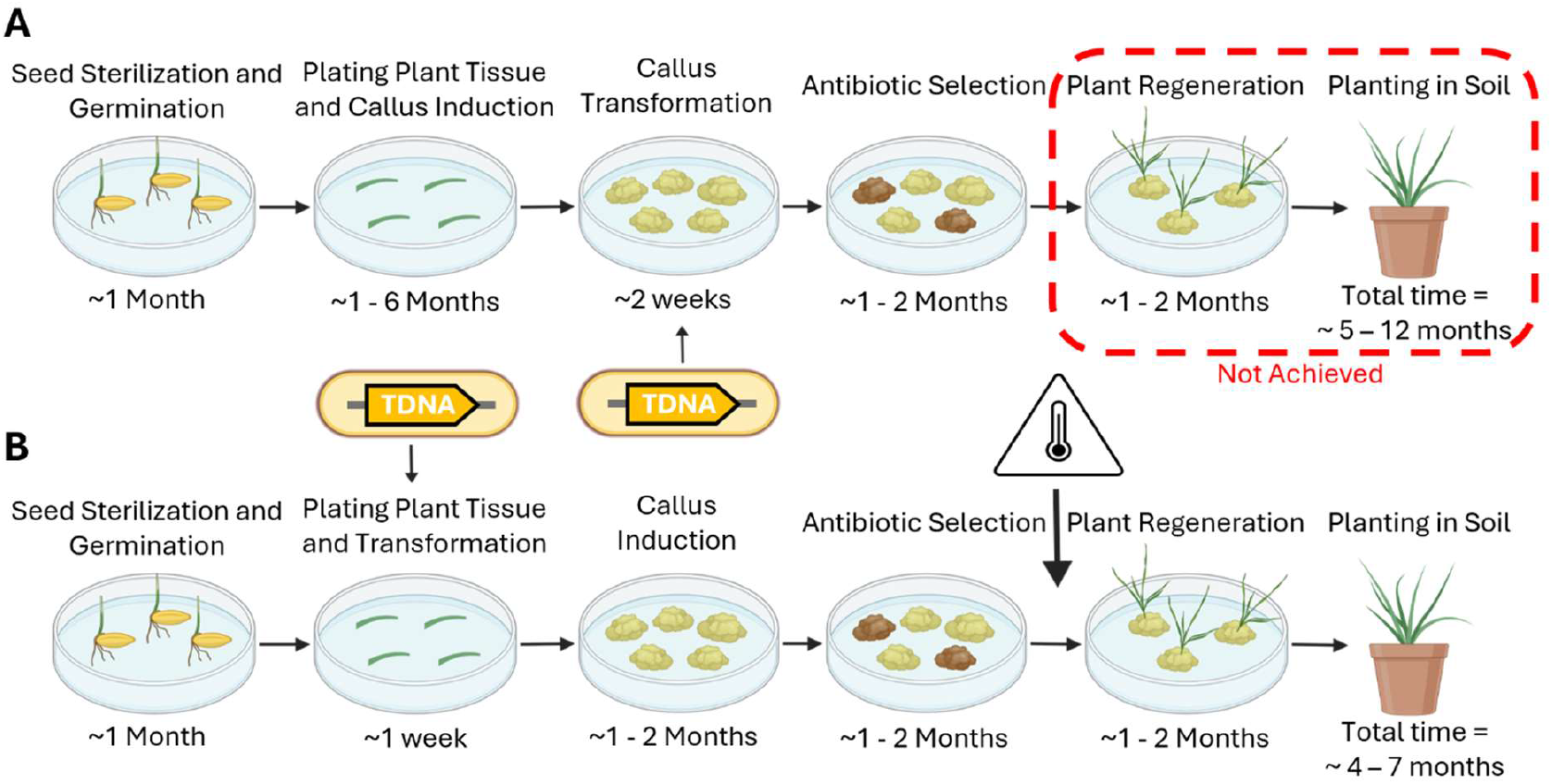
Schematic comparison of both protocols. Original callus protocol (A) in comparison to the new developmental regulator protocol (B). The original protocol was longer and unable to regenerate plants, while the new protocol is shorter and succeeded in creating transformed plants.

## Materials and Methods

### Key components

Culture components such as Murshag & Skroog medium; 2,4-D; BA; etc. were all purchased from Phytotech (Lenexa, KS). Enzymes were purchased from New England Biolabs (Ipswich,MA). Synthetic DNA such as the primers for PCR were purchased from Integrated DNA Technologies (Newark, NJ). Plasmid DNA was isolated utilizing Qiagen (Germantown, Maryland) QIAiqiuck spin columns. Perennial ryegrass was grown in Sungro (Agawam, MA) professional potting mix. Three perennial ryegrass cultivars were tested for callus formation: ‘SeaBiscuit’, ‘Pharaoh’, and ‘Arctic Green’. ‘Arctic Green’ was used for transformation.

### Protoplast methods

Protoplast isolation and transformations methods were adapted from Sychla et al. (2022) with key variables being taken from Yu et al. (2017) Briefly, 8 day old seedlings that had been growing in a mist house were finely cut-up and placed in enzyme solution. They were then vacuumed in the dark at 20 inHg for 30 minutes. The tissue was digested for 6 hours in a temperature-controlled room at 25°C. The protoplasts were then cleaned with two centrifugations at 100xg and re-suspensions in 5 ml W5 solution and finally purified with a 0.55 M sucrose gradient. A hemacytometer was used to measure the total amount of protoplasts. Protoplasts were then resuspended in MMG buffer to a concentration of 10^6^ protoplasts ml^-1^. Next, 10 ug of plasmid DNA, 500 ul of protoplasts, and 500 ul of 40% PEG4000 were mixed and let sit for 10 minutes in the dark. Protoplasts had 3.5 ml W5 added to them and then pelleted by centrifugation at 100xg for 5 minutes and washed with W5 solution. Protoplasts were allowed to rest overnight in W5 before readings were taken.

### Dual luciferase methods

Protoplasts were transformed with plasmids from Chamness et al. (2023) (Supplementary Table 1). After the protoplasts were allowed to rest overnight, they were lysed following instructions from the Dual-Luciferase Reporter Assay System by Promega (Madison, Wi). Luciferase fluorescence measurements were taken with a microplate reader (Glomax). Each biological replicate indicates a separate protoplast transformation. After an initial failed run which didn’t have any Renilla values, a secondary Renilla plasmid driven by the *35S* promoter was co-transformed to act as the transformation normalization factor.

### Media formulations

All media compositions named throughout are listed in Supplementary Table 2. For the preparation of all media, temperature stable components were mixed and adjusted to a pH of 5.7. The mixture was autoclaved, and the media containers were then cooled under a laminar flow hood. Once the media had cooled to touch, roughly 50°C, temperature sensitive components were added. Subsequently, 25 mL of the media was poured into each petri plate, cooled, sealed, and stored at 4°C until needed.

### Seed sterilization

Seeds of ‘Arctic Green’ were sterilized in a laminar flow hood following an adapted protocol from Ke and Lee (1996) with fungicide treatment taken from Bajaj et al. (2006). Seeds were first rinsed with 70% ethanol for 1 minute. The ethanol was then decanted using sterile pipettes and the seeds were rinsed 2-3 times with autoclaved water. Subsequently, 3% sodium hypochlorite (NaOCl) and 1% sodium dodecyl sulfate solution were added, and the seeds were placed on a shaker for 1 hour at approximately 100 rpm (light shaking). After decanting the NaOCl solution, the seeds were rinsed 2-3 times with autoclaved water. Next, 20 mL of 1.5% NaOCl and 1% SDS were added, and the seeds were shaken for an additional 30 minutes. Following decantation, the seeds were rinsed 5-6 times with autoclaved water. Approximately 5 mL of autoclaved water was added to the rinsed seeds, which were deposited onto sterile petri plates containing two pieces of autoclaved filter paper. The plates were sealed and placed at 10°C for 3-7 days of stratification.

### Sterile seed germination

After stratification, the seeds were removed from the 10°C storage and opened in a sterilized flow hood. The lightest color seeds were selected, and approximately 30 seeds were placed onto an agarose media plate containing fungicide (Supplementary Table 2). After all seeds were placed, the plates were sealed with micropore tape and incubated in a growth chamber at 21°C for two to four weeks.

### Original callus culture transformation methods

*Agrobacterium* cultures strain AGL1 were grown overnight in LB medium (10g/L tryptone, 5g/L yeast extract, 5g/L sodium chloride) that contained the appropriate antibiotics for the vector utilized (kanamycin), at 28°C and were shaken at 200 rpm. *Agrobacterium* suspensions were pelleted at 4,500 g for 8 minutes, and re-suspended in basal MS medium containing 1% (w/v) glucose and 3% (w/v) sucrose, 400 μM acetosyringone, pH 5.2, at an OD600 of 0.80. Approximately 30 calluses were then placed into 50 mL centrifuge tubes, or until they reached the 5 mL line. Calluses was fully immersed with the *Agrobacterium* liquid culture, shaken for 30 minutes at 100 rpm at room temperature, and then placed onto a petri plate with sterile filter paper to dry. Calluses were placed onto co-cultivation medium (Supplementary Table 2) for 4 days in the dark at 25°C and were then transferred to MS selection plates (Supplementary Table 2) with desired antibiotics (hygromycin) for 2 weeks at 25°C. Calluses were subcultured one time in the same medium with half the concentration of antibiotics. After one month, growing calluses were transferred to MS regeneration medium (Supplementary Table 2) with ¼ original concentration of antibiotics and sub-cultured every 2 weeks until plants were regenerated.

### Genetic construct creation

All constructs were cloned into TDNA backbones via the MoClo system outlined in Chamness et al. (2023) and utilizing the Golden Gate procedure (Engler et al., 2008). Full constructs are included in Supplementary Table 1.

### Developmental regulator methods

*Agrobacterium* strain LBA4404 cultures were streaked from glycerol stocks onto LB plates containing the antibiotics kanamycin and gentamicin at a concentration of 50 μg/ml. Four colonies were selected and inoculated into 50 mL of liquid LB containing 1 µL/mL kanamycin and gentamicin. The cultures grew at 28°C for two days. The OD600 of the cultures was measured, and the cultures were pelleted by centrifugation at 4,500xg for 8 minutes. The pellets were resuspended in infection medium and adjusted to an OD600 of 0.6.

Plantlets were transferred to sterile petri plates, and each plant was excised and placed into the infection medium. The explants were incubated on a shaker for 20 minutes at room temperature. They were then dried on autoclaved filter paper and transferred onto 710N co-cultivation medium (Supplementary Table 2). Sterile filter paper was placed on top of the medium to facilitate transfer. The plates were sealed with parafilm and incubated in the dark at 21°C for two days.

After co-cultivation, explants were transferred to resting media (Supplementary Table 2) for two weeks and subsequently transferred to selection media (Supplementary Table 2) containing kanamycin for two weeks at 21°C in the dark. To induce the heat shock response of the *Hsp17* promoter and activate the Cre recombinase, cultures were incubated at 45°C for two hours.

Explants were then transferred to maturation medium (404) (Supplementary Table 2) without filter paper and incubated at 28°C in the dark for two weeks. Afterward, the explants were moved to a 26°C light condition for one week. Transformed explants were transferred to the rooting medium (Supplementary Table 2) until regeneration or callus death occurred. Once regenerated roots and shoots were observed, they were transferred to soil and grown in the greenhouse.

### Transformation confirmation

After plants had been transferred to the greenhouse, DNA was extracted from a single leaf with DNAzol reagent following manufacturer’s instructions. PCRs utilizing Q5 DNA polymerase from NEB were conducted with the genomic DNA following manufacturer’s instructions and utilizing primers 5’-gctgctctagccaatacgca-3’ + 5’-acgagccggaagcataaagt-3’, and 5’-agcacagtggagtagggtat-3’ + 5’-attgtcaggttcggttctagtcttctt-3’ to confirm transgenesis and *cre* excision. PCR conditions were 98°C for 1 minute, 35 repeats of 98°C for 10 seconds, 68°C for 30 seconds, 72°C for 20 seconds, and a final extension at 72°C for 2 minutes. Conditions for *cre* excision were the same, except for the annealing temperature being 66°C and the extension time being 80 seconds.

## Data Availability

The data underlying this article are available in the article and in its online supplemental material.

## Funding

This work was supported by Hatch project award no. MN-21-096, from the U.S. Department of Agriculture’s National Institute of Food and Agriculture.

## Acknowledgments

We thank Dr. Dan Voytas and his lab members for support in curating and creating plasmid components. We thank Dr. Dominic Petrella for editing the manuscript.

## Author Contributions

JPC, MJS, and EW conceived this study. JPC and GWO designed and conducted experiments. JPC analyzed data. JPC, GWO, MJS & EW wrote the manuscript.

## Disclosures

### Conflicts of interest

No conflicts of interest declared

## Supplementary material

**Supplementary Figure 1.**
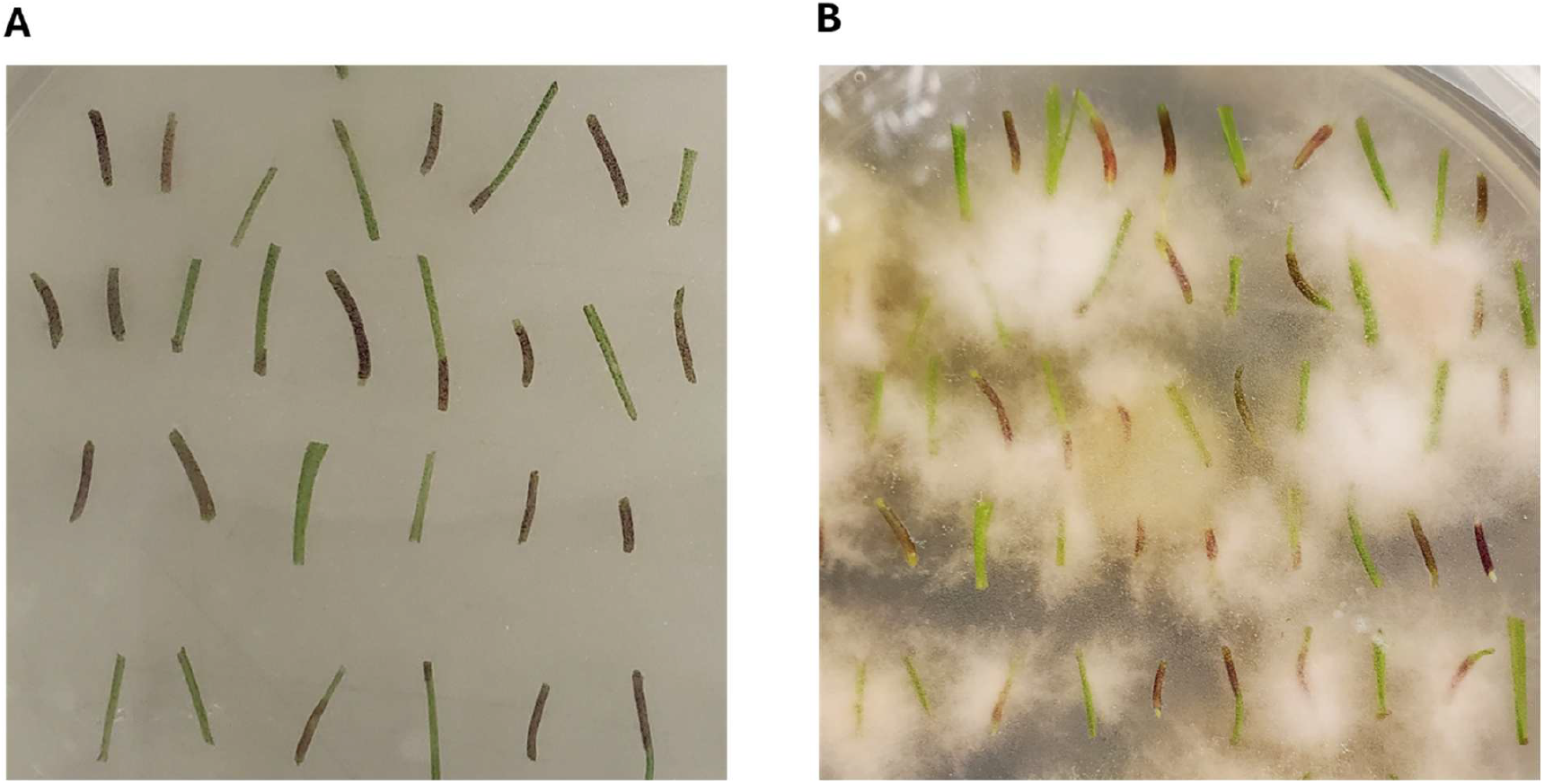
Before (A) and after (B) endophyte contamination of tissue on culture induction plates from tissue that wasn’t treated with fungicide.

**Supplementary Figure 2.**
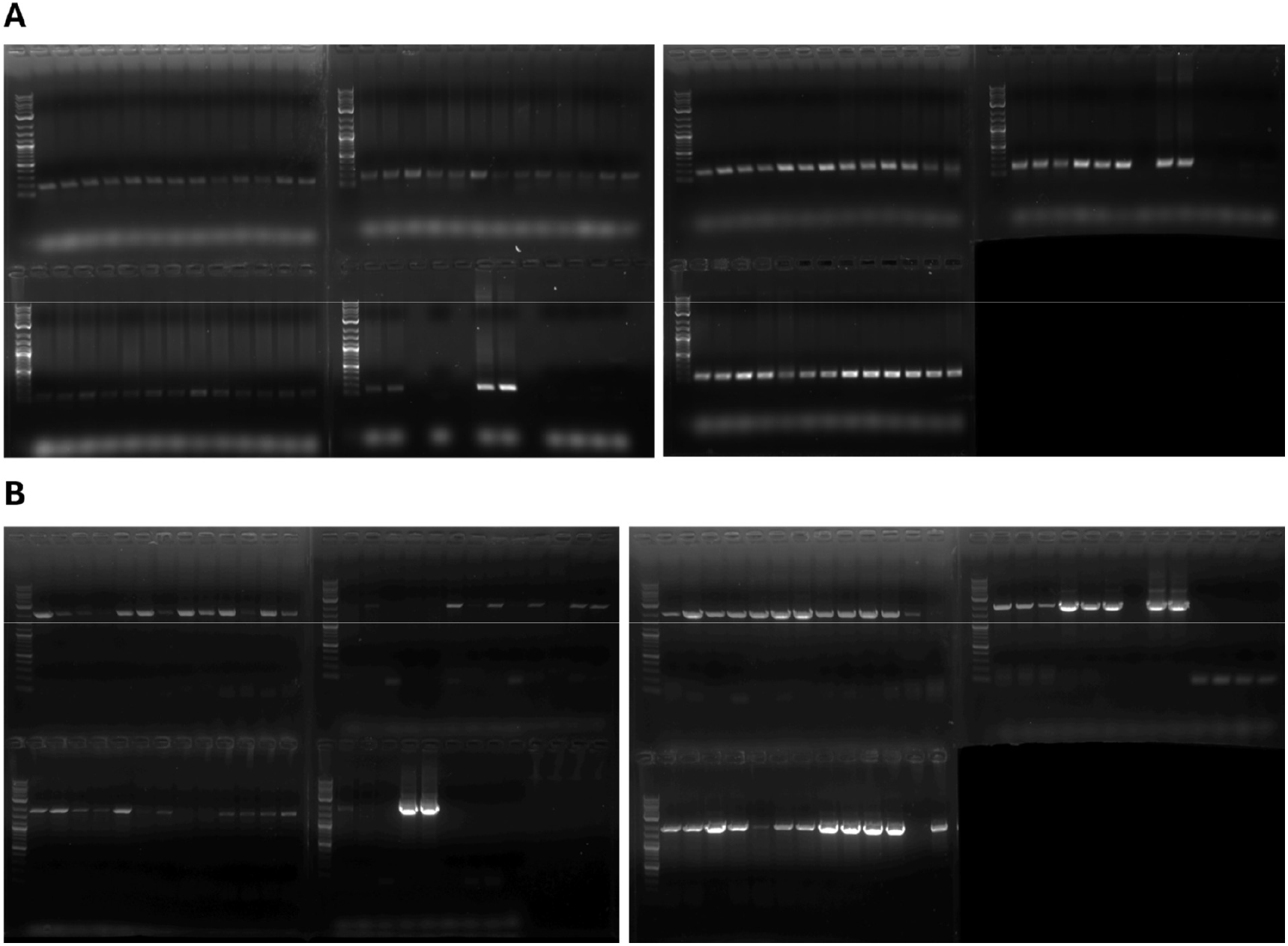
Full gels for confirmation of transformation (A) and cre excision (B). The first gel image is the first replicate, and the second gel is the second replicate. All wells before the final 6 are tested samples while the final 6 correspond to a no template control, Ubi/Zm/+ and nos/35/+ positive plasmid control, and 4 different wild type controls.

**Supplementary Figure 3.**
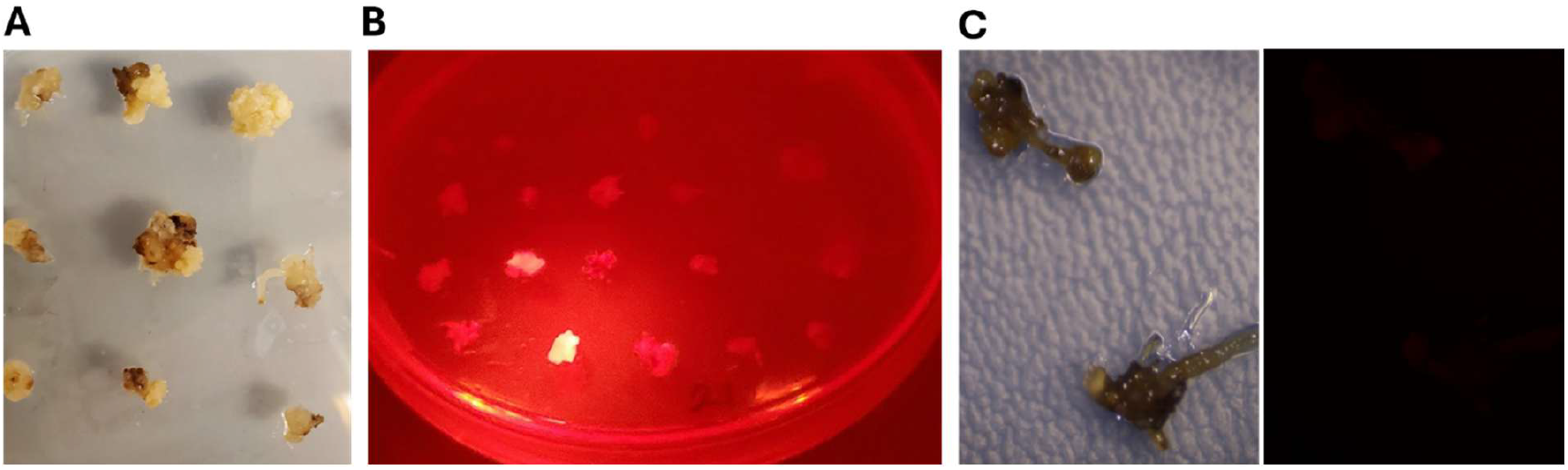
(A) Callus responding differently to antibiotic selection. (B) Differential expressions of RFP in callus. (C) Dead callus that didn’t survive antibiotic selection giving off no RFP signal.

**Supplementary Table 1.**
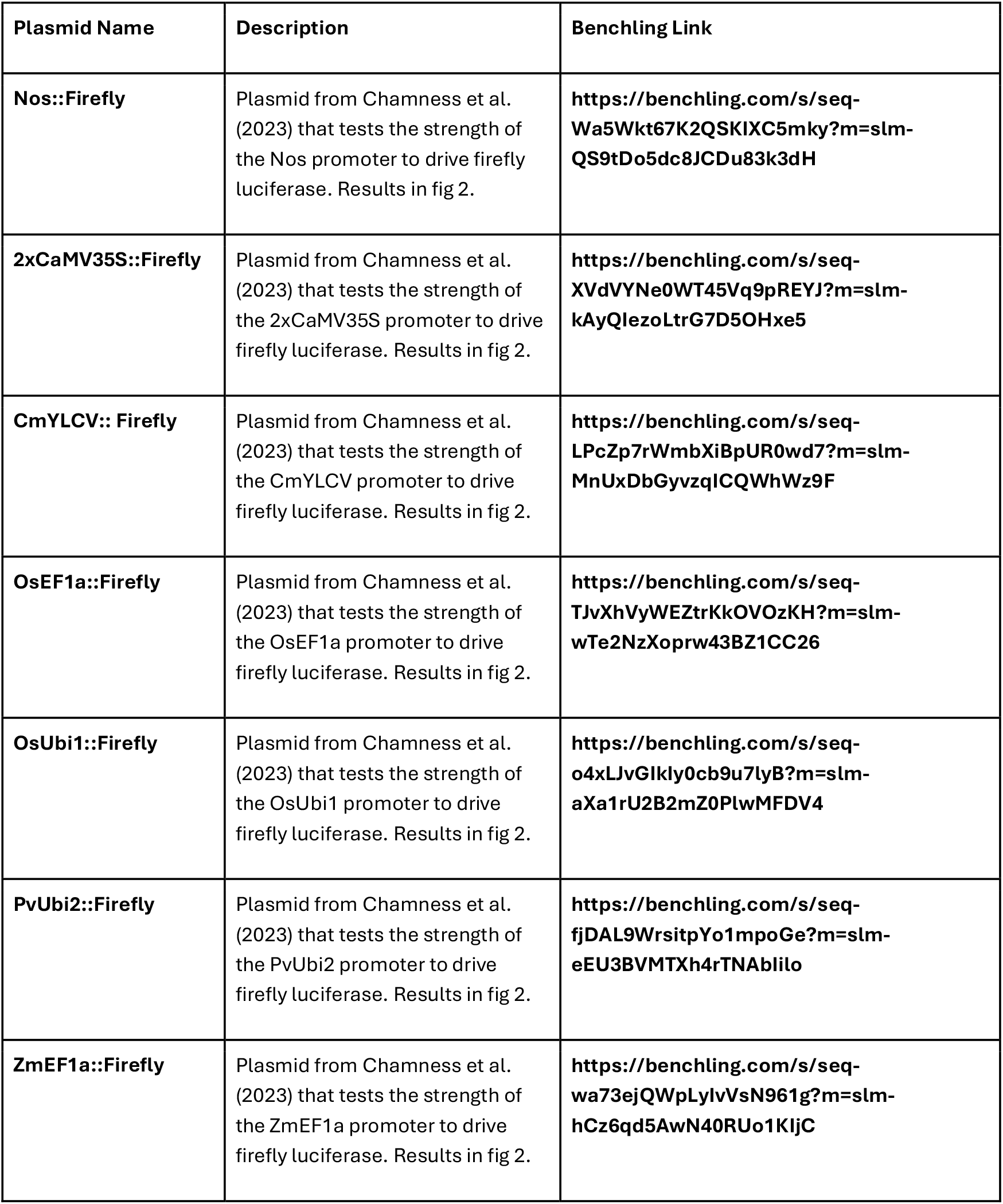
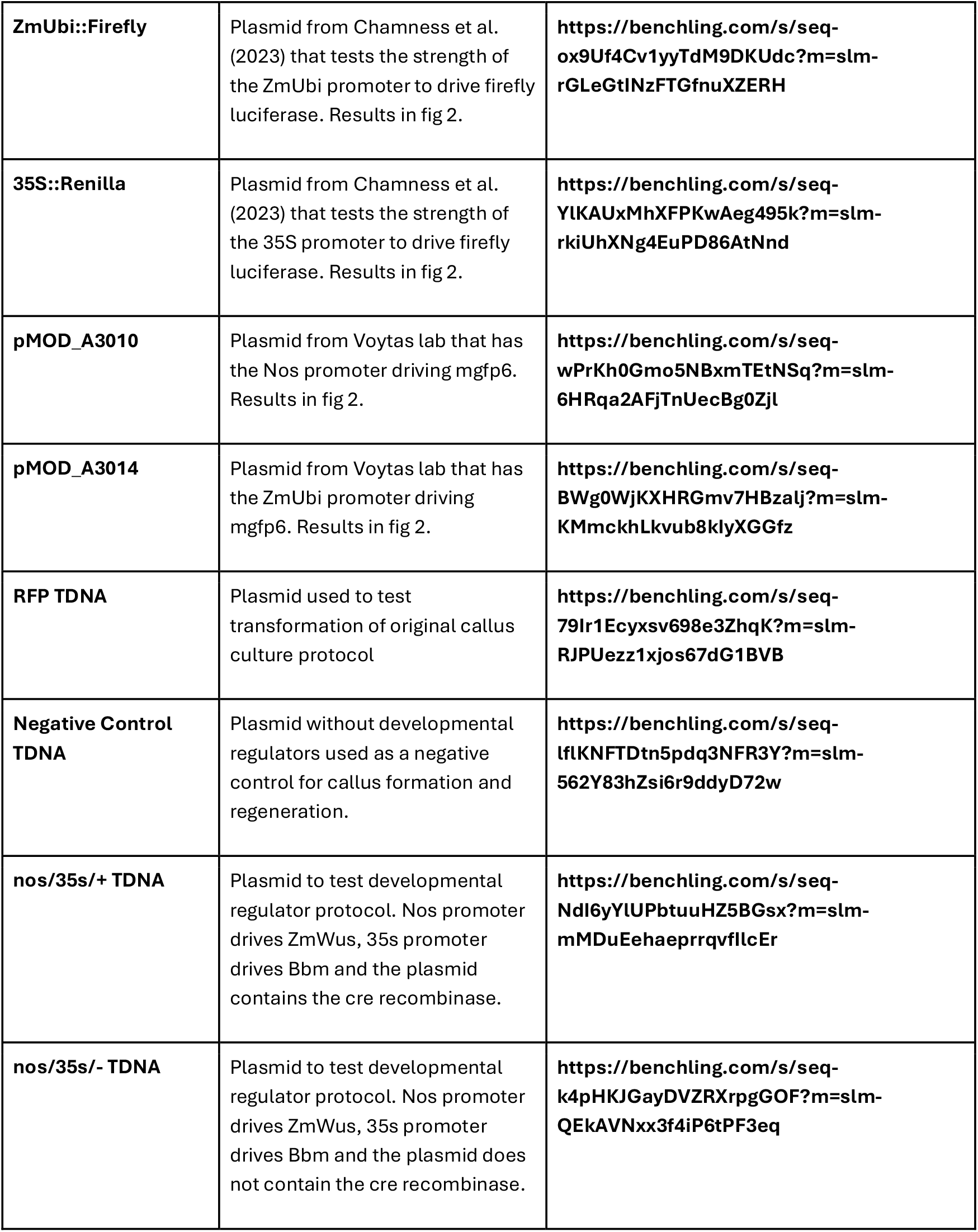
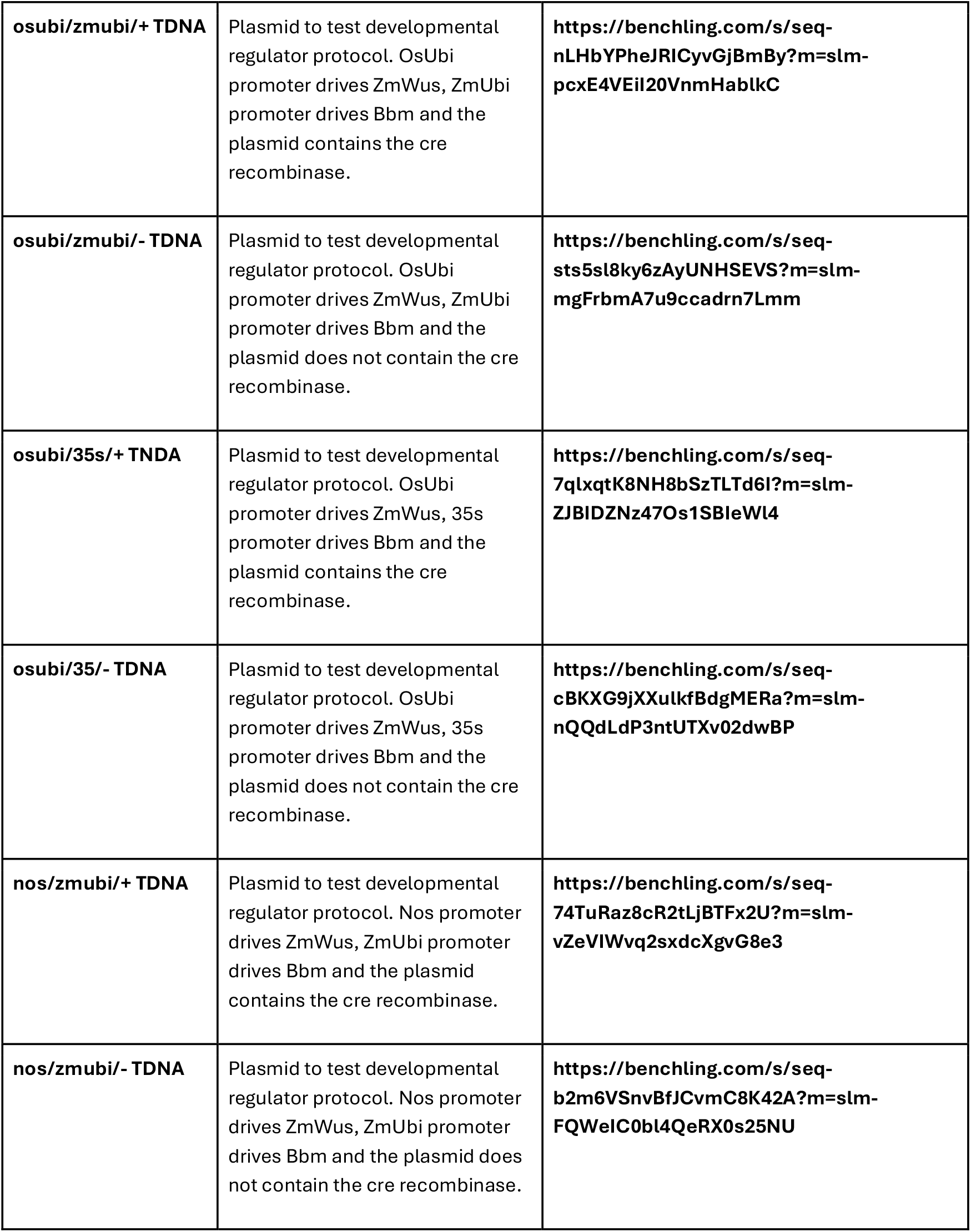
List of plasmids, their description, and benchling link to full plasmid map for every plasmid used in this paper.

**Supplementary Table 2.** List of every media and its contents used in the original callus culture protocol.

|  |  |
| --- | --- |
| **Fungicide Seed Plates** | 2.2g MS media for ½ strength<br>0.5g MES buffer<br>8g phytoblend for 0.8%<br>15g sucrose for 1.5%<br>10ml of 4g/L benomyl fungicide stock solution |
| **Culture Plates** | 4.4g MS media for full strength<br>0.5g MES buffer<br>8g phytoblend for 0.8%<br>30g sucrose for 3%<br>5ml of 1mg/ml 2,4-D stock solution |
| **Subculture Plates** | 4.4g MS media for full strength<br>0.5g MES buffer<br>8g phytoblend for 0.8%<br>30g sucrose for 3%<br>1ml of 0.1mg/ml BA stock solution<br>2ml of 1mg/ml 2,4-D stock solution |
| **Coculture Plates** | 4.4g MS media for full strength<br>0.5g MES buffer<br>8g phytoblend for 0.8%<br>30g sucrose for 3%<br>10g glucose for 1%<br>1ml of 0.1mg/ml BA stock solution<br>2ml of 1mg/ml 2,4-D stock solution<br>1ml of 400mM acetosyringone stock solution |
| **Subculture And Antibiotic Plates** | 4.4g MS media for full strength<br>0.5g MES buffer<br>8g phytoblend for 0.8%<br>30g sucrose for 3%<br>1ml of 0.1mg/ml BA stock solution<br>2ml of 1mg/ml 2,4-D stock solution<br>1ml of 50mg/ml hygromycin stock solution<br>250ul of 200mg/ml timentin stock solution |
| **Regeneration Plates** | 4.4g MS media for full strength<br>0.5g MES buffer<br>8g phytoblend for 0.8%<br>30g maltose for 3%<br>10ml of 0.1mg/ml BA stock solution<br>0.5ml of 50mg/ml hygromycin stock solution<br>250ul of 200mg/ml timentin stock solution |

**Supplementary Table 3.**
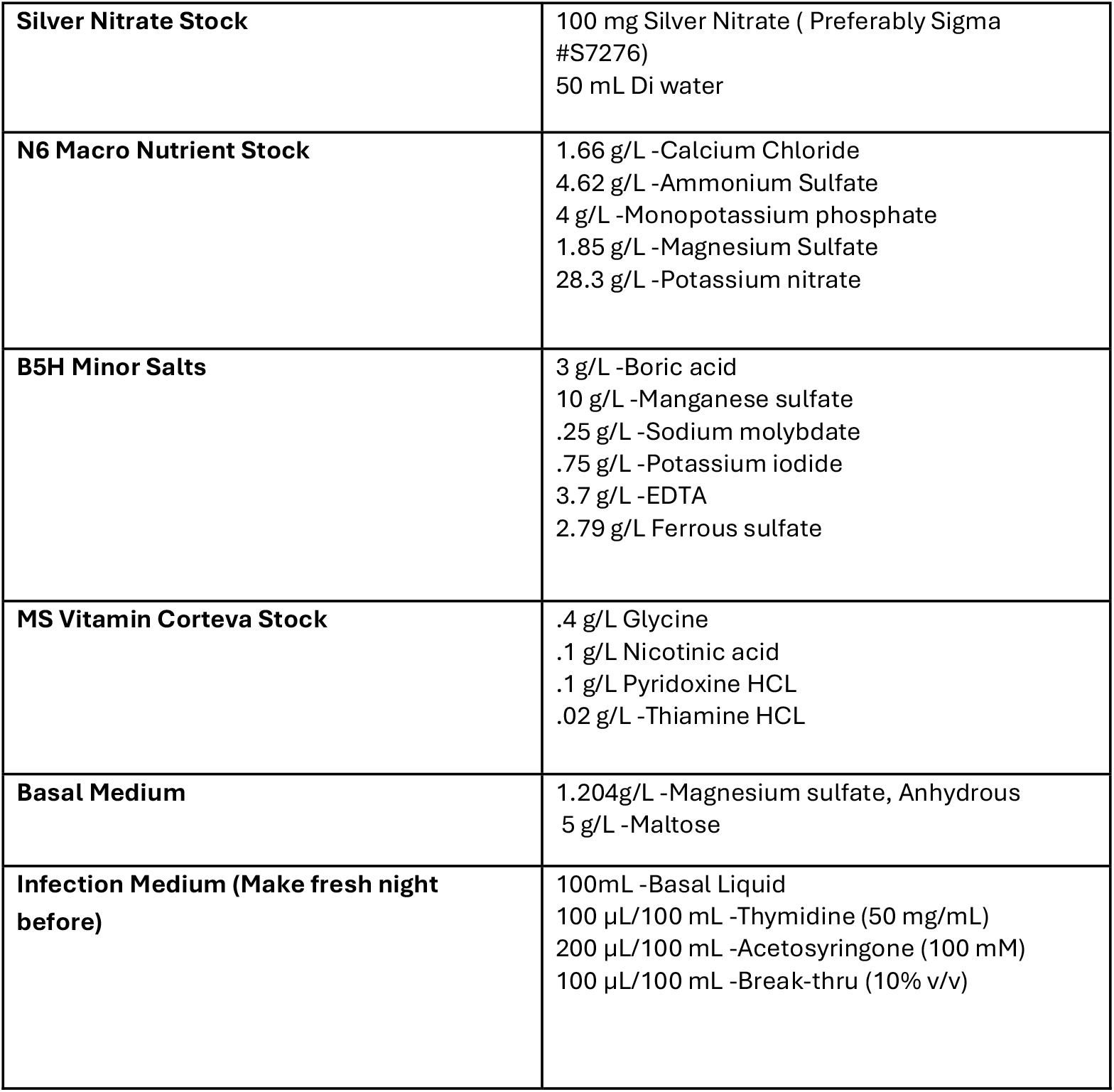
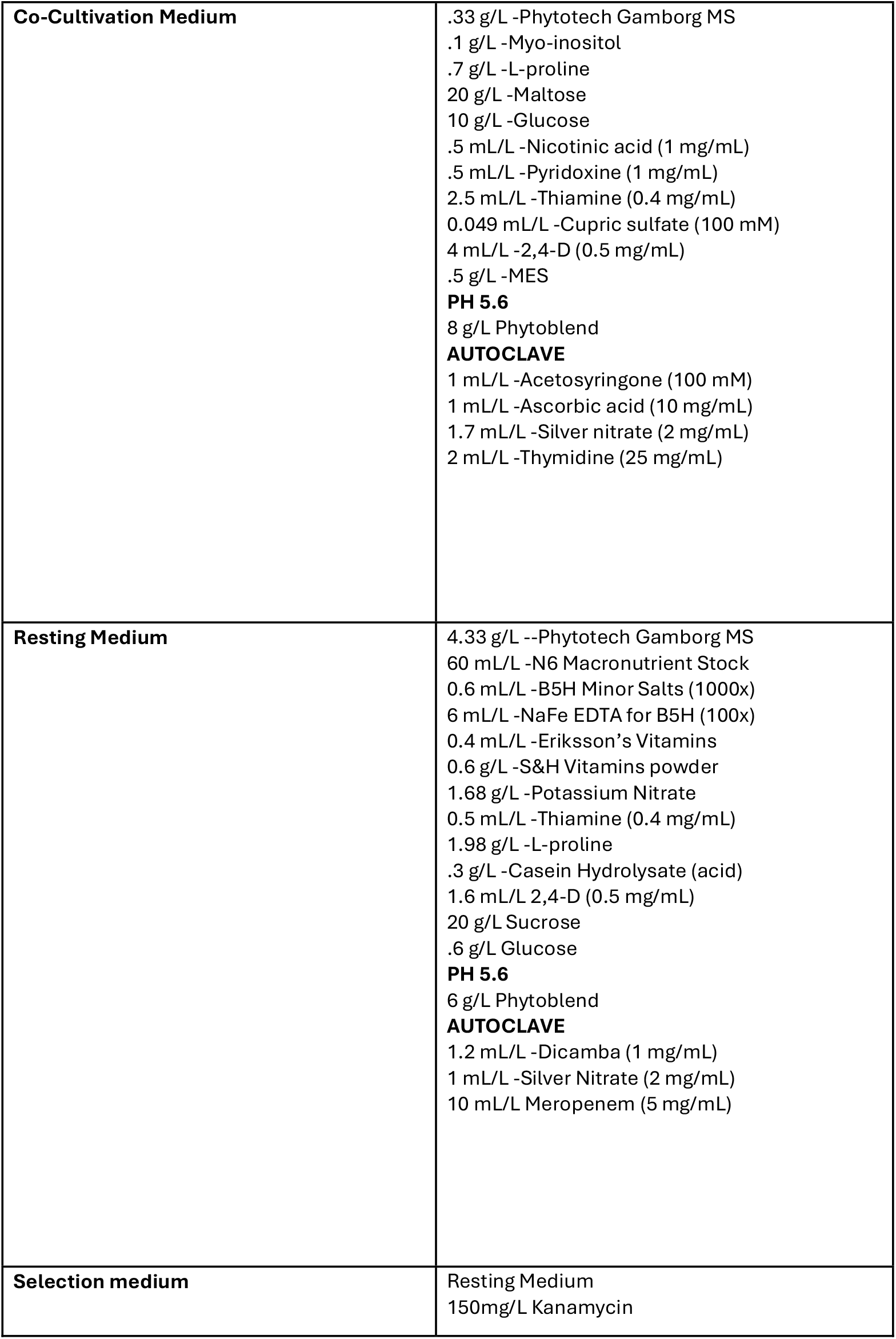
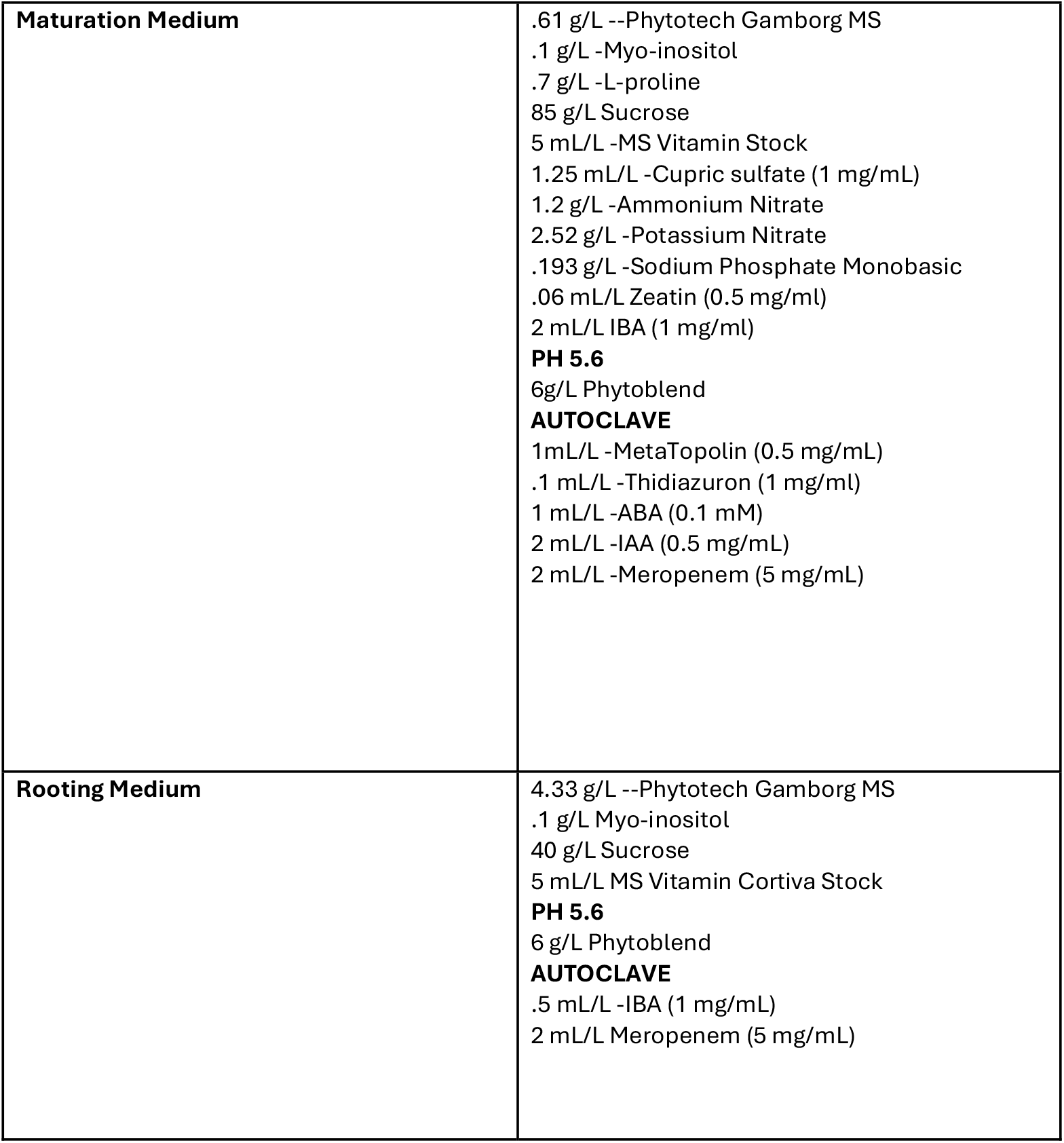
List of every media and its contents used in the developmental regulator protocol.

|  |  |
| --- | --- |
| **Silver Nitrate Stock** | 100 mg Silver Nitrate ( Preferably Sigma #S7276)<br>50 mL Di water |
| **N6 Macro Nutrient Stock** | 1.66 g/L -Calcium Chloride<br>4.62 g/L -Ammonium Sulfate<br>4 g/L -Monopotassium phosphate<br>1.85 g/L -Magnesium Sulfate<br>28.3 g/L -Potassium nitrate |
| **B5H Minor Salts** | 3 g/L -Boric acid<br>10 g/L -Manganese sulfate<br>.25 g/L -Sodium molybdate<br>.75 g/L -Potassium iodide<br>3.7 g/L -EDTA<br>2.79 g/L Ferrous sulfate |
| **MS Vitamin Corteva Stock** | .4 g/L Glycine<br>.1 g/L Nicotinic acid<br>.1 g/L Pyridoxine HCL<br>.02 g/L -Thiamine HCL |
| **Basal Medium** | 1.204g/L -Magnesium sulfate, Anhydrous<br>5 g/L -Maltose |
| **Infection Medium (Make fresh night before)** | 100mL -Basal Liquid<br>100 µL/100 mL -Thymidine (50 mg/mL)<br>200 µL/100 mL -Acetosyringone (100 mM)<br>100 µL/100 mL -Break-thru (10% v/v) |
| **Co-Cultivation Medium** | .33 g/L -Phytotech Gamborg MS<br>.1 g/L -Myo-inositol<br>.7 g/L -L-proline<br>20 g/L -Maltose<br>10 g/L -Glucose<br>.5 mL/L -Nicotinic acid (1 mg/mL)<br>.5 mL/L -Pyridoxine (1 mg/mL)<br>2.5 mL/L -Thiamine (0.4 mg/mL)<br>0.049 mL/L -Cupric sulfate (100 mM)<br>4 mL/L -2,4-D (0.5 mg/mL)<br>.5 g/L -MES<br>**PH 5.6**<br>8 g/L Phytoblend<br>**AUTOCLAVE**<br>1 mL/L -Acetosyringone (100 mM)<br>1 mL/L -Ascorbic acid (10 mg/mL)<br>1.7 mL/L -Silver nitrate (2 mg/mL)<br>2 mL/L -Thymidine (25 mg/mL) |
| **Resting Medium** | 4.33 g/L --Phytotech Gamborg MS<br>60 mL/L -N6 Macronutrient Stock<br>0.6 mL/L -B5H Minor Salts (1000x)<br>6 mL/L -NaFe EDTA for B5H (100x)<br>0.4 mL/L -Eriksson's Vitamins<br>0.6 g/L -S&H Vitamins powder<br>1.68 g/L -Potassium Nitrate<br>0.5 mL/L -Thiamine (0.4 mg/mL)<br>1.98 g/L -L-proline<br>.3 g/L -Casein Hydrolysate (acid)<br>1.6 mL/L 2,4-D (0.5 mg/mL)<br>20 g/L Sucrose<br>.6 g/L Glucose<br>**PH 5.6**<br>6 g/L Phytoblend<br>**AUTOCLAVE**<br>1.2 mL/L -Dicamba (1 mg/mL)<br>1 mL/L -Silver Nitrate (2 mg/mL)<br>10 mL/L Meropenem (5 mg/mL) |
| **Selection medium** | Resting Medium<br>150mg/L Kanamycin |
| **Maturation Medium** | .61 g/L --Phytotech Gamborg MS<br>.1 g/L -Myo-inositol<br>.7 g/L -L-proline<br>85 g/L Sucrose<br>5 mL/L -MS Vitamin Stock<br>1.25 mL/L -Cupric sulfate (1 mg/mL)<br>1.2 g/L -Ammonium Nitrate<br>2.52 g/L -Potassium Nitrate<br>.193 g/L -Sodium Phosphate Monobasic<br>.06 mL/L Zeatin (0.5 mg/ml)<br>2 mL/L IBA (1 mg/ml)<br>**PH 5.6**<br>6g/L Phytoblend<br>**AUTOCLAVE**<br>1mL/L -MetaTopolin (0.5 mg/mL)<br>.1 mL/L -Thidiazuron (1 mg/ml)<br>1 mL/L -ABA (0.1 mM)<br>2 mL/L -IAA (0.5 mg/mL)<br>2 mL/L -Meropenem (5 mg/mL) |
| **Rooting Medium** | 4.33 g/L --Phytotech Gamborg MS<br>.1 g/L Myo-inositol<br>40 g/L Sucrose<br>5 mL/L MS Vitamin Cortiva Stock<br>**PH 5.6**<br>6 g/L Phytoblend<br>**AUTOCLAVE**<br>.5 mL/L -IBA (1 mg/mL)<br>2 mL/L Meropenem (5 mg/mL) |

## Notes

### Competing Interest Statement

The authors have declared no competing interest.

## References

Bajaj, S., Ran, Y., Phillips, J., Kularajathevan, G., Pal, S., Cohen, D., et al. (2006) A high throughput Agrobacterium tumefaciens-mediated transformation method for functional genomics of Perennial ryegrass (Lolium perenne L.). Plant Cell Rep. 25: 651–659.

Bidabadi, S.S., and Jain, S.M. (2020) Cellular, molecular, and physiological aspects of in vitro plant regeneration. Plants. 9: 702.

Blume, C.J., Fei, S.-Z., and Christians, N.E. (2010) Field evaluation of reduced-growth, glyphosate-resistant Kentucky bluegrass in a noncompetitive setting. Crop Science. 50: 1048–1056.

Bonos, S.A., and Huff, D.R. (2015) Cool-season grasses: biology and breeding. In Turfgrass: Biology, Use, and Management. Edited by Stier, J.C., Horgan, B.P., and Bonos, S.A. pp. 591–660 American Society of Agronomy, Crop Science Society of America, Soil Science Society of America, Madison, WI, USA.

Braun, R.C., Mandal, P., Nwachukwu, E., and Stanton, A. (2024) The role of turfgrasses in environmental protection and their benefits to humans: Thirty years later. Crop Science. 64: 2909–2944.

Chamness, J.C., Kumar, J., Cruz, A.J., Rhuby, E., Holum, M.J., Cody, J.P., et al. (2023) An extensible vector toolkit and parts library for advanced engineering of plant genomes. The Plant Genome. 16: e20312.

Clark, M., and Maselko, M. (2020) Transgene biocontainment strategies for molecular farming. Front Plant Sci. 11.

Dai, W., Bonos, S., Guo, Z., Meyer, W., Day, P., and Belanger, F. (2003) Expression of pokeweed antiviral proteins in Creeping bentgrass. Plant Cell Rep. 21: 497–502.

Dündar, G., Ramirez, V.E., and Poppenberger, B. (2024) The heat shock response in plants: new insights into modes of perception, signaling, and the contribution of hormones. J Exp Bot. 76: 1970–1977.

Engler, C., Kandzia, R., and Marillonnet, S. (2008) A one pot, one step, precision cloning method with high throughput capability. PLoS One. 3: e3647.

Fan, J., Zhang, W., Amombo, E., Hu, L., Kjorven, J.O., and Chen, L. (2020) Mechanisms of environmental stress tolerance in turfgrass. Agronomy. 10: 522.

Gardner, D.S., Danneberger, T.K., Nelson, E., Meyer, W., and Plumley, K. (2003) Relative fitness of glyphosate-resistant Creeping bentgrass lines in Kentucky bluegrass. HortSci. 38: 455–459.

Gordon-Kamm, B., Sardesai, N., Arling, M., Lowe, K., Hoerster, G., Betts, S., et al. (2019) Using morphogenic genes to improve recovery and regeneration of transgenic plants. Plants. 8: 38.

Grogg, D., Rohner, M., Yates, S., Manzanares, C., Bull, S.E., Dalton, S., et al. (2022) Callus induction from diverse explants and genotypes enables robust transformation of Perennial ryegrass (Lolium perenne L.). Plants. 11: 2054.

Han, Y., Jin, X., Wu, F., and Zhang, G. (2011) Genotypic differences in callus induction and plant regeneration from mature embryos of barley (Hordeum vulgare L.). J Zhejiang Univ Sci B. 12: 399–407.

Hartman, C.L., Lee, L., Day, P.R., and Tumer, N.E. (1994) Herbicide resistant turfgrass (Agrostis palustris Huds.) by biolistic transformation. Nat Biotechnol. 12: 919–923.

Hu, B., Zhang, G., Liu, W., Shi, J., Wang, H., Qi, M., et al. (2017) Divergent regeneration-competent cells adopt a common mechanism for callus initiation in angiosperms. Regeneration (Oxf). 4: 132–139.

Ikeuchi, M., Sugimoto, K., and Iwase, A. (2013) Plant callus: mechanisms of induction and repression. The Plant Cell. 25: 3159–3173.

Ke, S., and Lee, C.W. (1996) Plant regeneration in Kentucky bluegrass (Poa pratensis L.) via coleoptile tissue cultures. Plant Cell Reports. 15: 882–887.

Kumar, R., Kamuda, T., Budhathoki, R., Tang, D., Yer, H., Zhao, Y., et al. (2022) Agrobacterium- and a single Cas9-sgRNA transcript system-mediated high efficiency gene editing in Perennial ryegrass. Front Genome Ed. 4.

Long, Y., Yang, Y., Pan, G., and Shen, Y. (2022) New insights into tissue culture plant-regeneration mechanisms. Front Plant Sci. 13.

Lowe, K., Wu, E., Wang, N., Hoerster, G., Hastings, C., Cho, M.-J., et al. (2016) Morphogenic regulators baby boom and wuschel Improve monocot transformation. The Plant Cell. 28: 1998–2015.

Luo, H., Kausch, A.P., Hu, Q., Nelson, K., Wipff, J.K., Fricker, C.C.R., et al. (2005) Controlling transgene escape in GM Creeping bentgrass. Mol Breeding. 16: 185–188.

Ma, Z., He, J., Shen, Y., Li, Y., Wang, P., and Duan, T. (2025) Impact of grass endophyte on leaf spot in Perennial ryegrass caused by Bipolaris sorokiniana and subsequent aphids’ feeding preference. Agriculture. 15: 116.

Maselko, M., Feltman, N., Upadhyay, A., Hayward, A., Das, S., Myslicki, N., et al. (2020) Engineering multiple species-like genetic incompatibilities in insects. Nat Commun. 11: 4468.

Petty, S., Yue, C., and Watkins, E. (2024) Investigating how nontariff measures impact the turfgrass seed trade. HortTechnology. 34: 412–423.

Reichman, J.R., Watrud, L.S., Lee, E.H., Burdick, C.A., Bollman, M.A., Storm, M.J., et al. (2006) Establishment of transgenic herbicide-resistant Creeping bentgrass (Agrostis stolonifera L.) in nonagronomic habitats. Molecular Ecology. 15: 4243–4255.

Santhanam, R., Luu, V.T., Weinhold, A., Goldberg, J., Oh, Y., and Baldwin, I.T. (2015) Native root-associated bacteria rescue a plant from a sudden-wilt disease that emerged during continuous cropping. Proceedings of the National Academy of Sciences. 112: E5013–E5020.

Singh, R.K., and Prasad, M. (2016) Advances in Agrobacterium tumefaciens-mediated genetic transformation of graminaceous crops. Protoplasma. 253: 691–707.

Sychla, A., Casas-Mollano, J.A., Zinselmeier, M.H., and Smanski, M. (2022) Characterization of programmable transcription activators in the model monocot Setaria viridis via protoplast transfection. In Protoplast Technology, Methods in Molecular Biology. Edited by Wang, K. and Zhang, F. pp. 223–244 Springer US, New York, NY.

Wang, J., Tan, M., Wang, X., Jia, L., Wang, M., Huang, A., et al. (2023) WUS-RELATED HOMEOBOX 14 boosts de novo plant shoot regeneration. Plant Physiol. 192: 748–752.

Wang, N., Ryan, L., Sardesai, N., Wu, E., Lenderts, B., Lowe, K., et al. (2023) Leaf transformation for efficient random integration and targeted genome modification in maize and sorghum. Nat Plants. 9: 255–270.

Wieners, R.R., Fei, S., and Johnson, R.C. (2006) Characterization of a USDA Kentucky bluegrass (Poa pratensis L.) core collection for reproductive mode and DNA content by flow cytometry. Genet Resour Crop Evol. 53: 1531–1541.

Wu, R., Chai, Y., Li, Y., Chen, T., Qi, W., Xue, Y., et al. (2025) A visual monitoring DNA-free multi-gene editing system excised via LoxP::FRT/FLP in poplar. Plant Biotechnol J. 23: 4017–4029.

Young, C.A., Hume, D.E., and McCulley, R.L. (2013) FORAGES AND PASTURES SYMPOSIUM: Fungal endophytes of tall fescue and perennial ryegrass: Pasture friend or foe? 12. J Anim Sci. 91: 2379–2394.

Yu, G., Cheng, Q., Xie, Z., Xu, B., Huang, B., and Zhao, B. (2017) An efficient protocol for perennial ryegrass mesophyll protoplast isolation and transformation, and its application on interaction study between LpNOL and LpNYC1. Plant Methods. 13: 46.

Zhang, Y., Ran, Y., Nagy, I., Lenk, I., Qiu, J.-L., Asp, T., et al. (2020) Targeted mutagenesis in ryegrass (Lolium spp.) using the CRISPR/Cas9 system. Plant Biotechnology Journal. 18: 1854–1856.

Zinselmeier, M.H., Casas-Mollano, J.A., Cors, J., Ferreira, S.S., Voytas, D.F., and Smanski, M.J. (2025) Towards engineering hybrid incompatibility in plants. Plant Biotechnology Journal. 23: 2752–2754.

